# Computational investigations in inhibition of alcohol/aldehyde dehydrogenase in lignocellulosic hydrolysates

**DOI:** 10.1101/2022.11.19.517192

**Authors:** Karan Kumar, Azeeza Siddiqa, Pragati, Priti Chandane, Saanya Yadav, Mahima Kori, A Shivram, Lepakshi Barbora, Vijayanand S. Moholkar

**Affiliations:** School of Energy Science and Engineering, Indian Institute of Technology Guwahati, Guwahati-781039, Assam, India.; Department of Biotechnology, Acharya Nagarjuna University, Guntur-522510, Andhra Pradesh, India.; Department of Botany, University of Delhi, University Road, Faculty of Science, University Enclave, Delhi-110007 India.; Department of Biochemistry, School of Life Sciences, University of Hyderabad, Hyderabad-500046, Telangana, India.; Department of Biotechnology, Delhi Technological University, Shahbad Daulatpur, New Delhi - 110042 Delhi NCR, India.; Department of Bioinformatics, Rajiv Gandhi Institute of IT and Biotechnology, Bharati Vidyapeeth, Pune, Maharashtra, India.; Department of Biochemistry, Pt. Jawahar Lal Nehru Memorial Medical College, Raipur, 492001, Chhattisgarh, India.; Department of Chemical Engineering, Indian Institute of Technology Guwahati, Guwahati-781039, Assam, India.

**Keywords:** Alcohol/Aldehyde dehydrogenase, Lignocellulosic biomass, Solventogenic Clostridia, Molecular Dynamic simulation, Density Functional Theory

## Abstract

Second generation alcoholic biofuels synthesis from lignocellulosic biomass (LB) consists three steps viz., pre-treatment, detoxification, and fermentation. This dilute acid pre-treatment process generates several compounds like acids, aldehydes, ketones, oxides and their phenolic derivatives that are potential inhibitors of some of the crucial enzymes in the metabolic pathway of ABE fermentation. With application of hybrid quantum mechanics/ molecular mechanics (QM/MM) approach, our aim is to discern the molecular mechanism of inhibition of key *AADs* across solventogenic species. The objectives of present study are: (1) identification and homology modelling of key *AADs*; (2) validation, quality assessment and physiochemical characterization of the modelled enzymes; (3) identification, construction and optimization of chemical structure of potent microbial inhibitors in LH; and (4) applications of hybrid QM/MM simulations to profile the molecular interactions between microbial inhibitors and key *AADs*. Our computational investigation has revealed various important facets of inhibition of the *AAD* enzymes, which could guide structural biologist in designing efficient and robust enzymes. Moreover, our methodology also provides a general framework which could applied for deciphering the molecular mechanism of inhibition behaviour of other enzymes.

**Highlights:**

- Homology modelling of 7 *alcohol/aldehyde dehydrogenase* (*AAD*) in solventogenic *Clostridia*
- Identification and structural optimization of potent microbial inhibitors in lignocellulosic hydrolysates
- QM/MM simulations to profile the molecular interactions between 10 inhibitors and 7 *AADs*
- Discernment of the molecular mechanism of inhibition of key *alcohol/aldehyde dehydrogenase*
- A methodological framework for deciphering the molecular mechanism of enzyme inhibition

## 1. Introduction

Second generation alcoholic biofuels make use of hydrolysates from lignocellulosic biomass as fermentation substrates. These hydrolysates are generated by the hydrolysis of the cellulosic and hemicellulosic fractions in biomass (Dahman, 2012; Kour et al., 2019; Mayank et al., 2013; Ranjan and Moholkar, 2012; Rathour et al., 2018). The hydrolysis of hemicellulosic fraction, which generate the hydrolysate rich in pentose sugars like xylose and arabinose, involves high temperature and pressure dilute acid treatment (Borah et al., 2016; Kumar et al., 2021). This process also generates several other organic compounds like acids, aldehydes, ketones, oxides and their phenolic derivatives, furan derivatives, and weak organic acids that are potential inhibitors of some of the crucial enzymes in the metabolic pathway (Agbor et al., 2011; Baral and Shah, 2014; Jönsson and Martín, 2016; Kim et al., 2011; Palmqvist and Hahn-Hägerdal, 2000). Bioethanol and biobutanol are potential alcoholic biofuels that can be blended with gasoline (Cao and Sheng, 2016; Kumar et al., 2022a, 2021; Malani et al., 2019; Singh et al., 2022). Both of these can be produced through acetone-butanol-ethanol (ABE) fermentation carried out by the *Clostridial* microbial cultures (Kolesinska et al., 2019; Kumar et al., 2022b; Mayank et al., 2013). The general metabolic pathway of ABE fermentation from pentose and hexose substrate is shown in Fig. 1(A). It could be seen that alcohol/ aldehyde dehydrogenase enzymes play a vital role in the formation of end-products (ethanol and butanol).

**Fig. 1.**
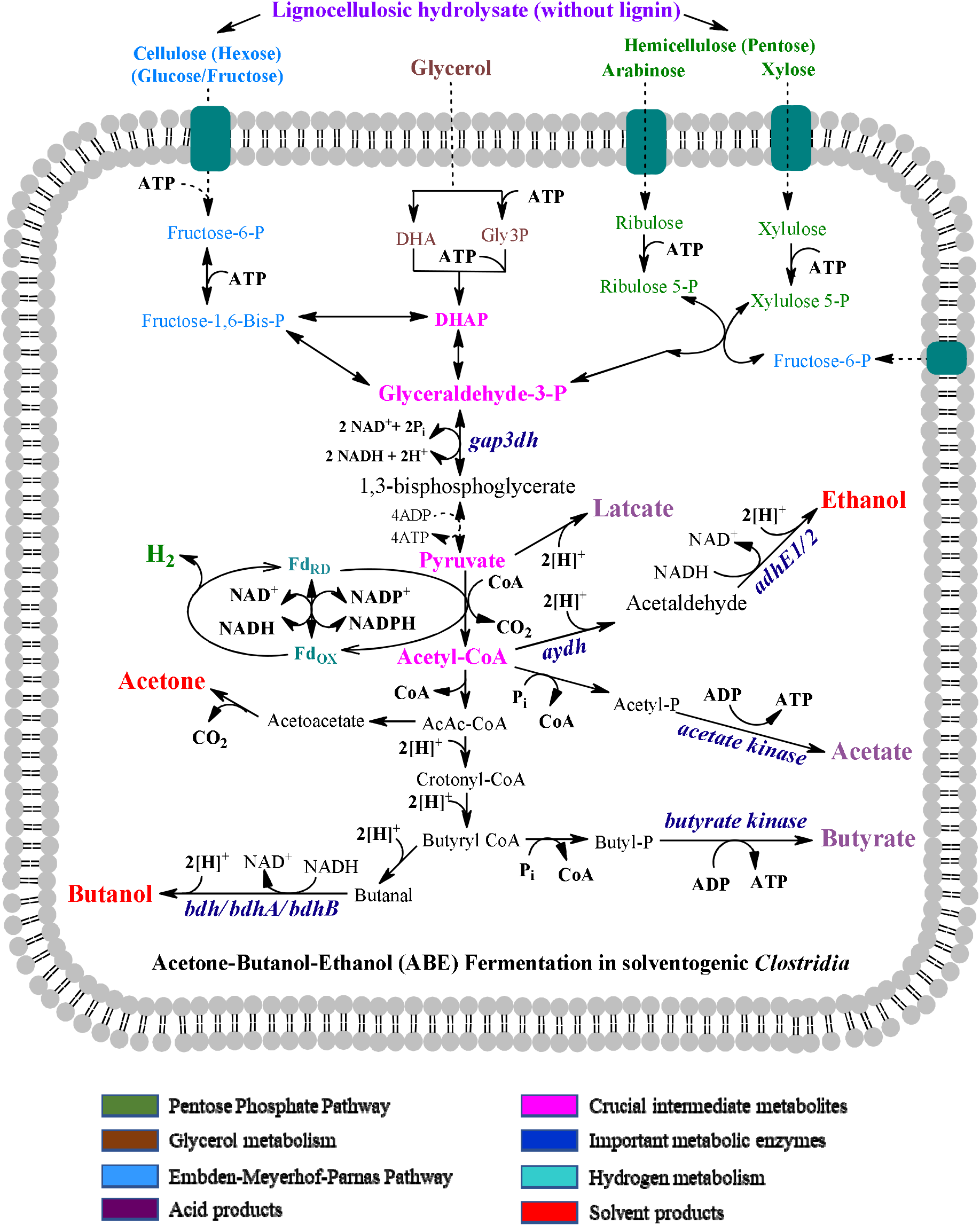
(A): Representative carbohydrate metabolism in solventogenic *Clostridia:* crucial solvent-producing enzymes, key metabolites, and brief acetone-butanol-ethanol (ABE) fermentation pathway.

As evident from the previous literature, *adhE1, adhE2, bdhA, bdhB, SMB P058*, and *yqhD* are the crucial *AAD* enzymes in metabolism of *C. acetobutylicum* leading to alcohols synthesis (Chen, 1995; Dai et al., 2016; Yoo et al., 2016). Similarly, the enzymes such as acetaldehyde dehydrogenase (*aydh*) and glyceraldehyde-3-phosphate dehydrogenase (*gap3dh*) have crucial role in metabolism of *Clostridium beijerinckii* and *Clostridium pasteurianum*, respectively. Fig. 1(A) reveals the exact role and location of these *AAD* enzymes in the cellular metabolism of ABE fermentation. From Fig. 1(A), it could be seen that *gap3dh* catalyse the conversion of glyceraldehyde-3-phosphate to 1,3-bisphosphoglycerate, an important step in the progression of glycolysis pathway. Solventogenic enzymes such as *aydh, adhE1, adhE2, bdh, bdhA, bdhB, acetate kinase*, and *butyrate kinase* play a vital role during ABE fermentation pathway. Bifunctional alcohol/ aldehyde dehydrogenase such *aydh* catalyses conversion of acetyl-coA to acetaldehyde, *adhE1* and *adhE2* catalyse conversion of an aldehyde (e.g. acetaldehyde or butanal) to alcohol (e.g. ethanol and butanol) and vice versa in ABE fermentation pathway. Similarly, the alcohol specific dehydrogenases such as *bdh*, *bdhA*, and *bdhB* also play important role in conversion of butanal to butanol, which is the final step of the butanol formation. The *AAD* enzymes are known to be inhibited by the aforementioned organic compounds present in the acid hydrolysate (Bhatia et al., 2020; Carrillo-Nieves et al., 2019; Fatma et al., 2018; Singhvi et al., 2014). This inhibition adversely affects not only the kinetics of metabolism but also the final yield of the ethanol and butanol products. With application of hybrid quantum mechanics/ molecular mechanics (QM/MM) approach, our aim is to discern the molecular mechanism of inhibition of key *AADs* across solventogenic species. The objectives of present study are: (1) identification and homology modelling of key *AADs;* (2) validation, quality assessment and physiochemical characterization of the modelled enzymes; (3) identification, construction and optimization of chemical structure of potent microbial inhibitors in LH; and (4) applications of hybrid QM/MM simulations to profile the molecular interactions between microbial inhibitors and key *AADs*. Our computational investigation has revealed various important facets of inhibition of the *AAD* enzymes, which could guide structural biologist in designing efficient and robust enzymes. Moreover, our methodology also provides a general framework which could applied for deciphering the molecular mechanism of inhibition behaviour of other enzymes.

## 2. Computational methodology

The schematic representation of workflow including general steps followed in this study is provided in Fig. 1B. Detailed procedure in step-by-step manner followed is presented in following sections.

**Fig. 1.**
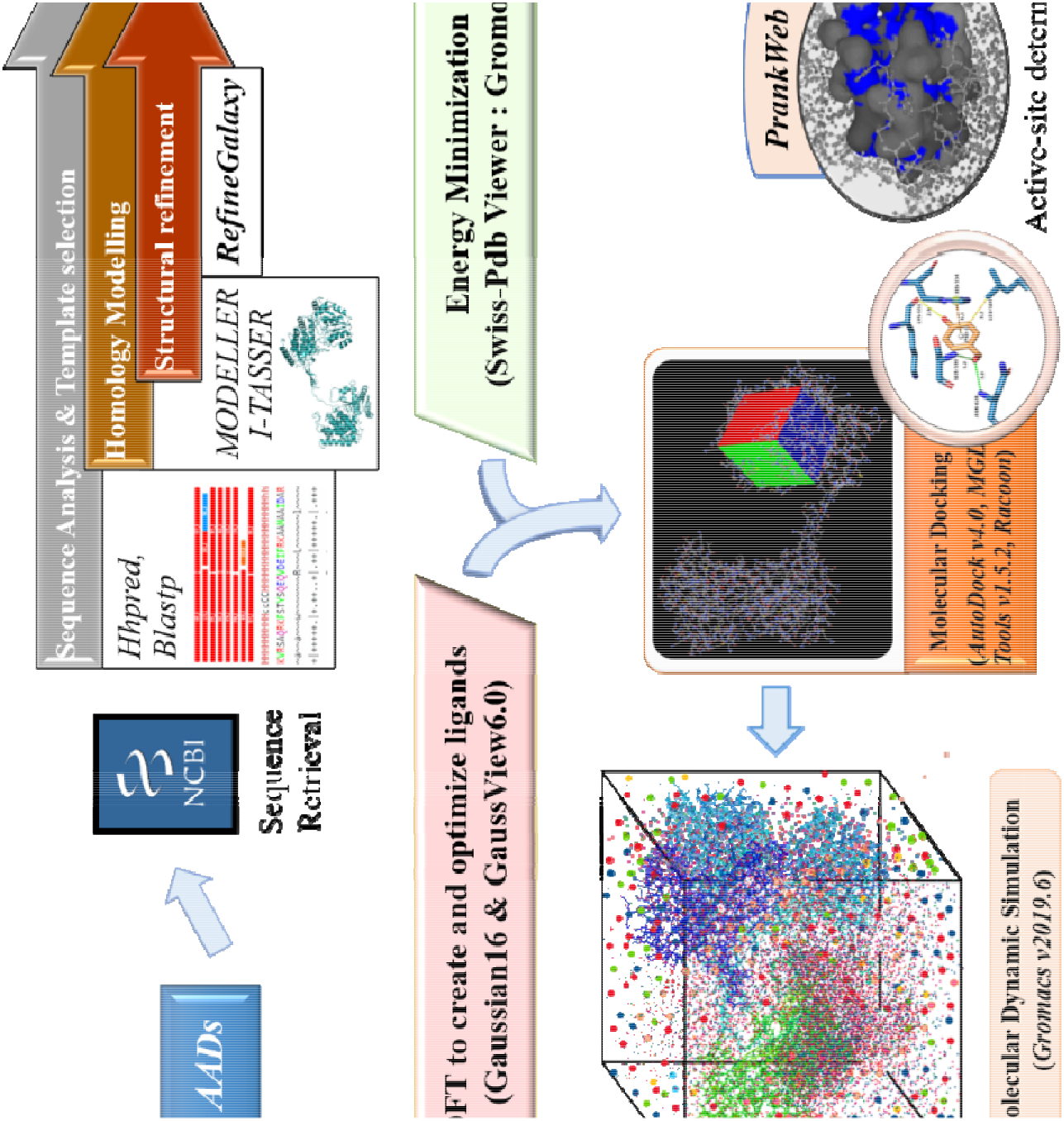
(B): Schematic workflow and methodology followed in the present study.

### 2.1. Preparation and analysis of key *AADs* in solventogenic *Clostridia*

Based on previous literature, we identified a total of seven *Clostridial AAD* enzymes (viz. *adhE1, adhE2, aydh, gap3dh, bdh, bdhA, and bdhB*), which are of high importance in solvent formation pathways (Chen, 1995; Cho et al., 2019; Dai et al., 2016). The metadata of these enzymes such as source organism, NCBI-refseq ID, sequence length, molecular weight and aliphatic index is listed in Table 1(A). The exact molecular functions and the structural informations, i.e. conserved domains, cofactors and conserved sites, is described in Table 1(B). In our study, we have classified *AAD* enzymes in four categories: (1) Bifunctional *AAD* enzymes that work on both alcohols and aldehydes (*adhE1* and *adhE2*), (2) butnol-specific *AAD* enzymes (*bdh, bdhA, and bdhB*), (3) biofunctional acetaldehyde dehydrogenase (*aydh*), and (4) glyceraldehyde metabolising dehydrogenase (*gap3dh*).

**Table 1.** (A): List of *Clostridial AAD* enzymes with their respective metadata from NCBI

| Enzyme name | Abb. | Source organism | Reference ID | # of aa | MW (kDa) | A.I. |
| --- | --- | --- | --- | --- | --- | --- |
| *Alcohol/aldehyde dehydrogenase enzyme 1* | *adhE1* | *Clostridium acetobutylicum* ATCC824 | AAK76781.1 | 858 | 94.44 | 92.65 |
| *Alcohol/aldehyde dehydrogenase enzyme 2* | *adhE2* | *Clostridium acetobutylicum* ATCC824 | AAK09379.1 | 858 | 94.44 | 92.65 |
| *NAD(P)H-dependent butanol dehydrogenase A* | *bdhA* | *Clostridium acetobutylicum* ATCC824 | OOM03255.1 | 389 | 42.98 | 91.26 |
| *NAD(P)H-dependent butanol dehydrogenase B* | *bdhB* | *Clostridium acetobutylicum* ATCC824 | OOM03254.1 | 390 | 43.29 | 97.26 |
| *Butanol dehydrogenase* | *bdh* | *Clostridium acetobutylicum* ATCC824 | AAA23206.1 | 389 | 43.07 | 91.00 |
| *Acetaldehyde dehydrogenase* | *aydh* | *Clostridium beijerinckii* NCIMB8052 | WP_026886587 | 864 | 94.93 | 91.53 |
| *Glyceraldehyde-3-phosphate dehydrogenase* | *gap3dh* | *Clostridium pasteurianum* ATCC6013 | OMH20402.1 | 334 | 36.08 | 92.54 |

Abbreviation: Abb. – abbreviation; aa – amino acids; MW – molecular weight; A.I. – aliphatic index;

**Table 1.** (B): Selected *Clostridial AAD* enzymes: their functions and other structural information

| Enzyme | Molecular functions | Conserved domains (residues range) | Cofactors | Conserved site |
| --- | --- | --- | --- | --- |
| *adhE1/E2* | • oxidoreductase activity<br>• metal ion binding<br>• alcohol dehydrogenase (NAD<sup>+</sup>) activity<br>• oxidoreductase activity, acting on the aldehyde or oxo group of donors, NAD or NADP as acceptor<br>• butanol dehydrogenase activity<br>• acetaldehyde dehydrogenase (acetylating) activity | Aldehyde dehydrogenase (5-374)<br>Iron-type/glycerol dehydrogenase (454-846)<br>Alcohol dehydrogenase (454-846)<br>Bifunctional aldehyde-alcohol dehydrogenase, C-terminal (453-855) | Zn<sup>2+</sup>, Fe<sup>2+</sup>,<br>NAD<sup>+</sup>, NADP<sup>+</sup> | 630 – 658<br>718 – 738 |
| *bdh/A/B* | • metal ion binding<br>• oxidoreductase activity<br>• butanol dehydrogenase activity | Iron containing dehydrogenase (9-381) | Zn<sup>2+</sup>, Fe<sup>2+</sup> | 173 – 201<br>264 – 284 |
| *Aydh* | • oxidoreductase activity<br>• oxidoreductase activity, acting on the aldehyde or oxo group of donors, NAD or NADP as acceptor<br>• metal ion binding<br>• acetaldehyde dehydrogenase (acetylating) activity<br>• alcohol dehydrogenase (NAD<sup>+</sup>) activity<br>• butanol dehydrogenase activity | Aldehyde dehydrogenase (5-398)<br>Iron-type/glycerol dehydrogenase (456-847)<br>Alcohol dehydrogenase (454-856)<br>Bifunctional aldehyde-alcohol dehydrogenase, C-terminal (454-856) | Zn<sup>2+</sup>, Fe<sup>2+</sup>, NAD<sup>+</sup>,<br>NADP<sup>+</sup> | 631-659<br>719-739 |
| *gap3dh* | • oxidoreductase activity, acting on the aldehyde or oxo group of donors, NAD or NADP as acceptor<br>• NAD binding<br>• NADP binding | NAD(P)-binding (2-150)<br>NAD-binding (2-102)<br>Catalytic (155-312)<br>C-terminal (155-312) | NAD<sup>+</sup>, NADP<sup>+</sup> |  |

#### 2.1.1. Sequence retrieval and secondary structure analysis

The amino acid sequences in FASTA format for all seven *AADs* were retrieved from the NCBI-RefSeq database. Selected metadata information for all seven *AADs* are provided in Table 1(A). Next, Protparam webtool (https://www.expasy.org/resources/protparam) was used for physicochemical analysis of retrieved protein sequence. Parameters computed using webtool Protparam in the query amino acid sequence are: (1) theoretical molecular weight, (2) molar extinction coefficient, (3) atomic composition, (4) theoretical pI (isoelectric point), (5) number of amino acids (AA), (6) composition of AA, (7) positively charged residues (Asp+Glu), (8) negatively charged residues (Arg+Lys), (9) aliphatic index, (10) grand average of hydropathicity (GRAVY), (11) theoretical half-life, and (12) instability index.

#### 2.1.2. Homology modelling of AAD enzymes

The 3-dimentional (3D) models of *AADs* were generated using MODELLER (Gabler et al., 2020) and I-TASSER (Roy et al., 2010). In case of MODELLER, templates for homology modelling were selected using HHpred (Zimmermann et al., 2018), which is inbuilt package of Bioinformatics Toolkit. The HHpred (http://toolkit.tuebingen.mpg.de/hhpred) selects template structures based on Hidden Markov Models’ (HMM) pairwise profile comparison. Apart from HHpred, BLASTp against PDB database was also used to predict the templates for homology modelling. Final templates were selected based on best score and query coverage for each enzyme. In contrast to MODELLER, I-TASSER webserver requires query sequence in FASTA format as input for the generating 3D models. I-TASSER (https://zhanglab.ccmb.med.umich.edu/I-TASSER/) builds 3D models of any protein based on *ab-initio* modelling approach (Beg et al., 2018). The output 3D models of I-TASSER and MODELLER were compared and model having highest confidence score (C-score) were selected. C-score is a quantitative estimate for the superiority of the predicted models by I-TASSER which typically ranges from −5 to 2. A higher c-score value signifies a model with high confidence and vice versa. List of templates used for homology modelling of *AADs* and their metadata are provided in Table 2(A).

**Table 2.**
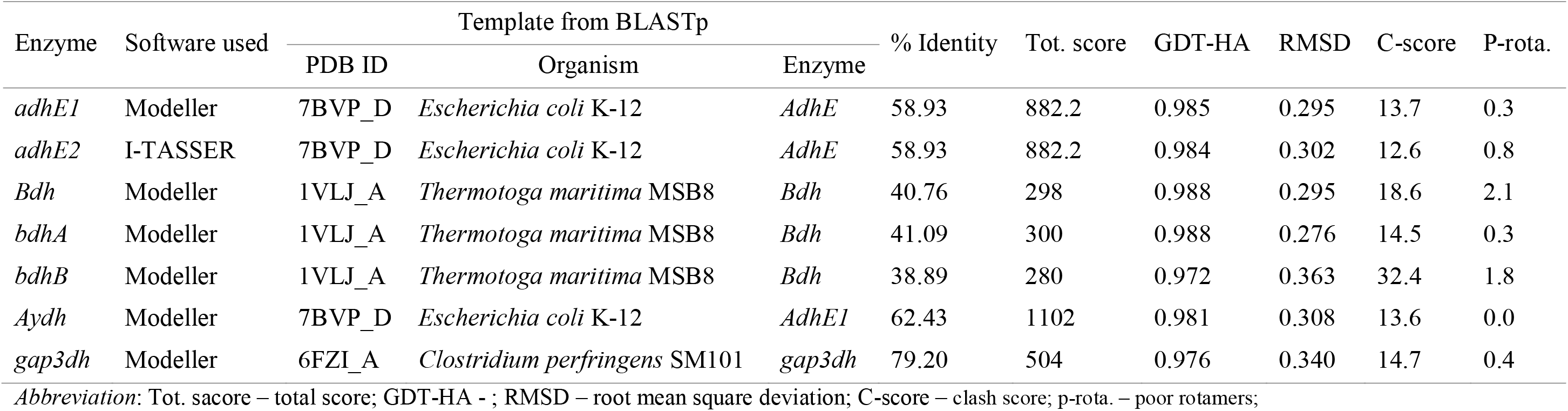
(A): List of templates used for homology modelling of *Clostridial AAD* enzymes and their metadata

#### 2.1.3. Refinement and validation of 3D structures of AADs

The selected enzyme models in the previous step were subjected further for model refining in GalaxyRefine (http://galaxy.seoklab.org/cgi-bin/submit.cgi?type=REFINE) to improve the quality. GalaxyRefine is an online server based on a CASP10 refining process that have been proven effective for model refining (Heo et al., 2013). GalaxyRefine uses small molecular dynamics simulations to reconstruct all side-chain conformations and relax the structure repeatedly. This method improves both global and local structure quality as determined by GDT-HA and root mean square deviation (RMSD). In the next step, Molprobity was used to evaluate the quality of backbone structure of final 3D models from GalaxyRefine. After the steps of structural refinement and energy minimization, the best 3D models (selected on the basis of Ramachandran score) were validated using various 3D structural parameters. Model validations were performed using Structural Analysis and Verification Server (SAVES v6.0: https://saves.mbi.ucla.edu/). Ramachandran plot analysis were carried out using PROCHECK to access the quality of final models. The stereochemical and geometrical parameters of the model were checked using the PROCHECK (Laskowski et al., 1996). Moreover, ERRAT server was used to examine the quality of the final 3D structure by determining the compatibility of an atomic model (3D) with its amino acid sequence and comparing the findings to standard structures (Wallner, 2006). After validation step, CASP14 (Critical Assessment of Structure Prediction) ranked webservers such as PrankWeb (available on P2Rank; Jendele et al., 2019), InterPro, 2Struc, and NetSurfP3.0 were used for identifying the ligand-binding pockets, putative catalytic residues, putative active site residues, putative metal-binding site, co-factor binding residues, secondary structural composition, and percentage surface accessibility analysis.

### 2.2. Selection of inhibitors and their preparation for hybrid QM/MM simulations

As noted earlier, the dilute acid pretreatment of LB produces many organic compounds, such as aliphatic/aromatic acids, aldehydes, furans, which act as metabolic and growth inhibitors and reduce the kinetics and yield of ABE fermentation. The toxic compounds also strongly inhibit the growth of vital cell structures and the metabolic capacity of fermenting strains.

#### 2.2.1. Identification and selection of inhibitors and representative substrates of AADs

Among various organic compounds produced during acid hydrolysis of LB, 10 compounds were identified as inhibitors of bacterial cell growth and enzymes in solventogenic pathway of metabolism (Baral and Shah, 2014; Kim, 2018; Yang et al., 2018). These inhibitors are 4-hydroxy-benzaldehyde (4-HBZ), benzaldehyde (BZD), cinnamaldehyde (CMD), syringaldehyde (SGN), ferulic acid (FA), levulinic acid (LA), vanillic acid (VA), *ρ*-coumaric acid (*ρ*-CA), vanillin (VN), and 5-hyroxymethylfurfural (HMF or 5-furfural). In addition, the representative substrates were identified for each *AAD* as follows: methanol (MET), ethanol (ETH), butanol (BUT), formaldehyde (FMD), acetaldehyde (ACD), butyraldehyde (BLD), and glyceraldehyde-3-phosphate (G-3-P). Hereon, the inhibitory compounds are referred as “fermentative inhibitors (FI)”. Further, the FIs in combination with representative substrates (as mentioned above) will be termed as “ligands”.

#### 2.2.2. Designing the structures of ligands and optimizing their geometry using Density Functional Theory (DFT)

The chemical structures of aforementioned ligands were designed in GaussView v6.0 software. Drawn structures in GaussView are not energetically favorable, hence, energy minimization is critical for establishing the optimal molecule arrangement in 3D space. The 3D geometry of ligands was optimized using DFT at B3PW91/6-311G(+)(d, p) level of theory followed by frequency calculation (freq = raman) in Gaussian16 software. The molecule’s potential energy components such as stretching, bending, and torsion, all of which are important in defining its behavior were optimized using Gaussian16 (Curtiss et al., 2007; “Gaussian 09 Citation | Gaussian.com,” n.d.). Finding the energy minima (i.e., the potential energy hypersurface of a stable molecule) helped improve the structure’s strain energy and, in turn, lead to the most stable conformer. The most stable conformer of the ligands was converted in *“.pdbqt”* format for further usage.

### 2.3. Molecular docking simulation and analysis

Prior to performing molecular docking, 3D model of *AAD* enzymes were subjected for energy minimization using SWISS-PDB Viewer (SPDBV; Guex and Peitsch, 1997), to achieve local minima that are closer to the native structure. The molecular docking simulations were performed to investigate biomolecular interactions, bound conformations, binding free energy of ligands and macromolecules, and identification of the active site residues using Autodock linked with the MGLTools version 1.5.6. The docking simulations of 17 ligands (combinations of 10 FIs and 7 representative substrates) against seven *AADs* were performed using Autodock v1.5.6 (Morris et al., 2009). The format of ligand structures generated through Gaussian16 were converted into *“.pdbqt”* format using OpenBabel software. The procedure for molecular docking was followed as described in Kumar et al. (2022a, 2021). The polar hydrogen ions as well as charges such as Kollman, Gasteiger were added to neutralize the system. A grid box was formed pertaining to coordinates for binding pockets using PrankWeb server. 100 docked conformations were generated using the Lamarckian Genetic Algorithm for investigating the optimal binding site of ligands in the *AAD* enzymes. For investigating various types of interactions, viz. polar and non-polar, in protein-ligand complexes various visualization software were used which include Autodock (Morris et al., 2009, 2008), PyMOL™ v2.4.1 (Yuan et al., 2017), PLIP (Protein-Ligand Interaction Profiler; https://plip-tool.biotec.tu-dresden.de/plip-web/), and VMD. LigPlot+ v.2.2 (Laskowski and Swindells, 2011) and Biovia’s Discovery studio was used to generate the 2D schematic representation of the interactions in enzyme-ligand complex. This protocol was reiterated for every ligand, and final optimal conformation possessing the lowest binding energy (among the hundred conformations procured for every operation) was analyzed carefully.

### 2.4. Molecular dynamic (MD) simulations of selected docked complexes of ligands with AADs

The MD simulations were performed using GROMACS v2019.3. The Charmm36-July2021 force field was employed in all MD simulations. The ligand topologies obtained after molecular docking for all ligands were submitted to the webserver PRODRG to generate GROMACS readable CHARMM coordinates. The obtained topologies (in “.*gro*” format) were integrated with respective *AAD* enzyme to form ligand-bound complexes. All the complexes were then solvated with water using the SPC/E water model. Adequate amount of chlorine or sodium ions were added to obtain the neutralized system. Next, the total energy of the system was minimized using the steepest descent method and then the system was equilibrated at 310 K and 1 atm by using NVT and NPT ensemble, respectively. This equilibrated system was submitted for production run for 50 ns each. As a conventional approach the PBC (periodic boundary conditions) were applied in all directions. Three simulation sets for each system were generated and data are represented as an average over the three sets to check the reproducibility and statistical significance of the results. The final MD trajectories was then analyzed in standalone VMD software. This methodology was repeated for each MD-simulated complex.

### 2.5. Analysis of MD-trajectories

After the MD simulations, trajectories were analysed and various parameters were calculated using different packages defined within GROMACS v.2019.3. The global structural stability of the inhibitor-bound complexes was determined by Cα-RMSD (Root Mean Square Deviation) by “*gmx rms*” package. The radius of gyration (Rg) for each complex were calculated using “*gmx gyrate*” package. Root mean square fluctuation (RMSF) and solvent-accessible surface area (SASA) have also been calculated to deduce detailed structural changes in the *AADs* by FI. All the results given are an average of three computational sets. Most potent inhibitors that destabilize the *AADs* were further accessed by enumerating the post-binding changes in secondary structure content of respective *AADs* by “*gmx do_dssp*” package. The hydrogen bonds play an important role in stabilization of any enzyme or protein, therefore, the destruction of hydrogen bonds was determined by the “*gmx hbond*” package. The intra-chain hydrophobic contacts for different residue pairs were calculated for each *AAD* in presence of ligands.

## 3. Results and discussions

### 3.1. Sequence and homology modelling of *AADs*

The FASTA sequences for all seven *AADs* were retrieved from NCBI-RefSeq database and analyzed as described in section 2.1.1. Template predicted by HHpred and BLASTp of query *AADs* sequences against PDB database showed similarity with previously characterized dehydrogenases. The *AADs* from bifunctional alcohol/aldehyde dehydrogenase category (i.e. *adhE1* and *adhE2*) and bifunctional acetaldehyde dehydrogenase (*aydh*) had highest percentage identity with Bifunctional aldehyde-alcohol dehydrogenase *AdhE* from *Escherichia coli* K12 (PDB id: 7BVP “chain D”) having 58.93% and 62.43%, respectively. All *AADs* from alcohol specific dehydrogenase categories (i.e. *bdh*, *bdhA*, and *bdhB*) had highest percentage identity with butanol dehydrogenase from *thermotoga maritima* MSB8 (PDB ID: 1VLJ “chain A”) with 40.76%, 41.09%, and 38.89%, respectively. However, *gap3dh* from aldehyde specific dehydrogenase category had highest percentage similarity with glyceraldehyde phosphate dehydrogenase from *Clostridium perfringens* SM101 (PDB ID: 6FZI “chain A”) with 79.20%. The crystal structures of 7BVP_D, 1VLJ_A, and 6FZI_A in PDB format were retrieved from RCSB-PDB database and used as template for homology modelling of respective *AADs* enzymes in MODELLER. Table 2B summarize the results from BLASTp template prediction and homology modelling simulations for each *AADs*. Standalone MODELLER software integrated with MPI toolkit and I-TASSER webserver were used to build the 3D structures of *AAD* enzymes.

**Table 2.** (B): Structural evaluation, validation, and secondary structural analysis of *Clostridial AAD* enzymes

| Enzymes | Ramachandran plot statistics (in %) |  |  |  | Errat score | Secondary structural composition (in %) |  |  | Surface accessibility (in %) |  |
| --- | --- | --- | --- | --- | --- | --- | --- | --- | --- | --- |
| | FR | AR | GAR | DR | | α-helices | β-sheets | Random coil | Exposed residue | Buried residue |
| *adhE1* | 94.2 | 5.3 | 0.1 | 0.4 | 91.4 | 49.5 | 13.2 | 37.3 | 42.3 | 57.7 |
| *adhE2* | 92.3 | 6.6 | 0.5 | 0.6 | 95.8 | 48.7 | 12.8 | 38.5 | 42.3 | 57.7 |
| *bdhA* | 95.3 | 4.1 | 0.6 | 0.0 | 94.1 | 54.8 | 10.8 | 34.4 | 43.2 | 56.8 |
| *bdhB* | 91.9 | 6.7 | 1.2 | 0.3 | 93.8 | 51.3 | 10.5 | 38.2 | 41.5 | 58.5 |
| *Bdh* | 97.9 | 1.8 | 0.3 | 0.0 | 96.3 | 55.5 | 10.3 | 34.2 | 43.2 | 56.8 |
| *Aydh* | 94.6 | 5.0 | 0.0 | 0.4 | 92.3 | 47.6 | 12.4 | 40.0 | 43.3 | 56.7 |
| *gap3dh* | 93.2 | 6.1 | 0.3 | 0.3 | 91.6 | 28.8 | 28.8 | 42.3 | 47.3 | 52.7 |

Abbreviation: FR - favoured region; AR - allowed region; GAR – generously allowed region; DR - disallowed region

### 3.2. Refinement and validation of modelled structure

The backbone ψ and Φ dihedral angles of the amino acid residues of all 7 *AADs* were analysed by Ramchandran plot through PROCHECK server. The homology modelled structures through I-TASSER and MODELLER showed anomaly (most of 3D models had Ramachandran scores less than 75%) which were again refined by GalazyRefine webserver. The final 3D models selected after refining were of improved quality and had more than 90% residues in favoured region of the Ramachandran plot and less than 1% of residues in disallowed region. The overall stereochemistry of each residue in the refined structures were checked using the Ramachandran plot. The results of Ramachandran analysis for final 3D models are summarised in Table 2B. Further, the quality assessments of final 3D models of all 7 *AADs* were performed using UCLA SAVESv.6.0 server, where the structures were mentioned pass by VERIFY 3D as residues ≥ 95.7% had an average 3D-1D score ≥ 0.2, which indicated the compatibility of amino acids in the modelled structure. Analysis of overall quality factor of the modelled 3D structures of *AAD* enzymes were performed by ERRAT server which was found to be ≥ 91% for each structure, which further established the excellent quality of the final 3D models of *AADs* (Table 2B). The goodness factor (g-factor) is a measure for identifying whether a stereo chemistry characteristic is “normal” or “abnormal.” The g-factor for *adhE1 adhE2, bdh, bdhA, bdhB, aydh*, and *gap3dh* were 0.19, 0.16, 0.15, 0.16, 0.03, 0.18, and 0.11, respectively. The acceptable g-factor values in PROCHECK are in the range of 0 to 0.5, with the best models having g-factor values close to zero, suggesting that the model is of high quality. All 3D models in this study falls in this range suggesting our models were of excellent quality. The Fig. 2A depicts the final 3D structure of *AAD* enzymes visualized in PyMOL. These 3D models showed α/β/α type fold.

**Fig. 2.**
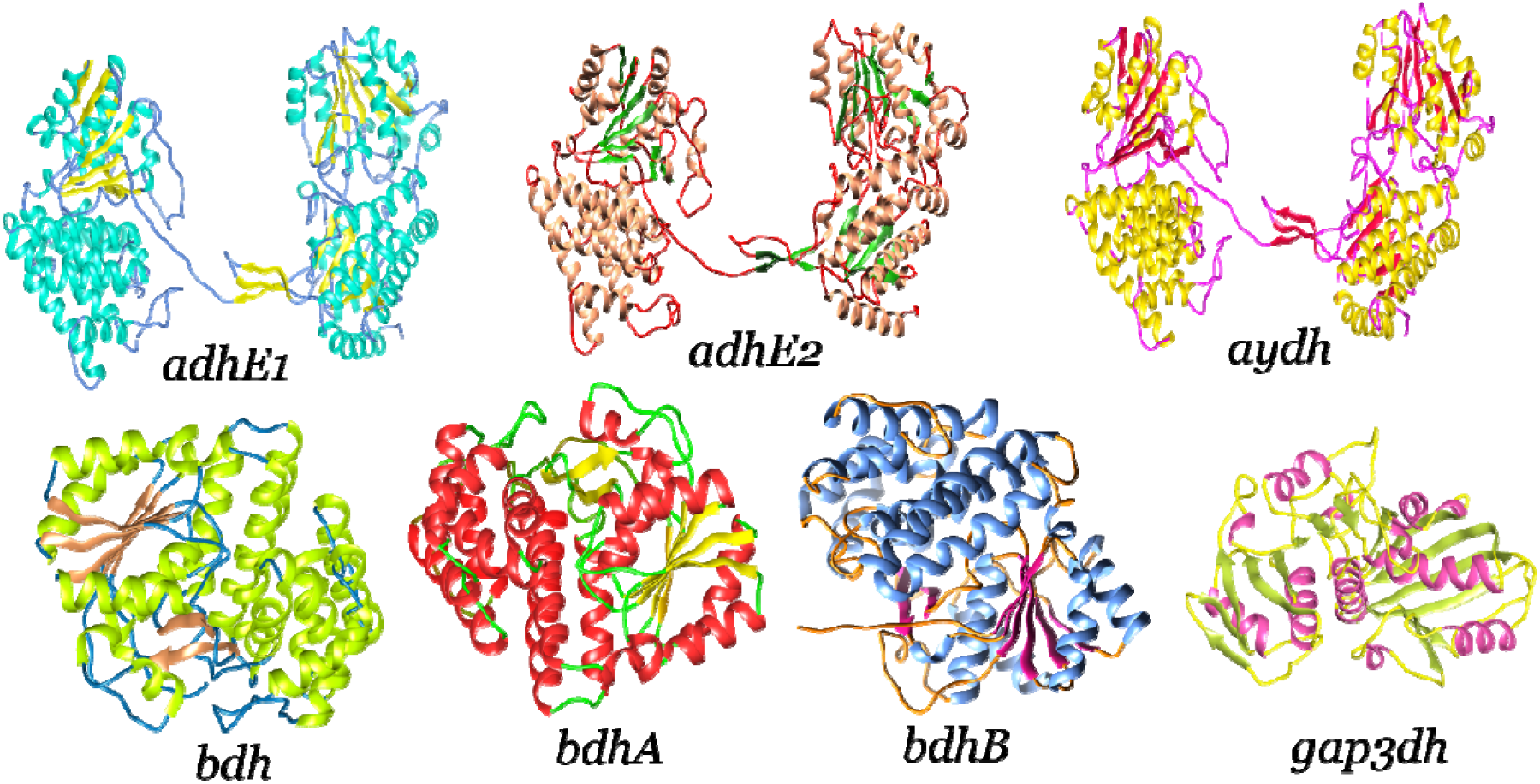
(A): Final 3D structure of 7 *Clostridial AAD* enzymes after homology modelling, refinement, and validation.

### 3.3. Secondary structure and binding pocket analysis of modelled *AADs*

Post modelling and validation of the final models, each enzyme was subjected to various CASP14 ranked webservers for secondary structural and binding pocket analysis. These CASP14-ranked webservers include PrankWeb, InterPro, 2Struc, 2StructCompare, PsiPred, and NetSurfP3.0 for identifying the ligand-binding pockets, putative catalytic residues, putative active site residues, putative metal-binding site, co-factor binding residues, secondary structural composition, and percentage surface accessibility (buried and exposed residues). PrankWeb that utilize machine learning algorithm to annotate the protein structure, was used for ligand-binding site prediction. In the process of predicting ligand-binding sites, points with a high predicted ligandability are grouped and scored using a ranking process based on the cluster’s cumulative score. The secondary structural analysis performed through 2Struc and PsiPred along with surface accessibilities are provided in Table 2B. All *AAD* enzymes except *gap3dh*, has higher α-helical content (approximately 48 to 56%) with approximately 73 to 76% surface accessibility (percentage ratio of exposed to buried residues) of the enzymes. The *gap3dh* enzyme has higher random coil content (42.3%) with approximately 90% surface accessibility. The prediction of catalytic residue showed presence of nucleophilic cysteine, serine or histidine residues in each *AAD* enzyme. The residues present in active site, metal-binding site, cofactor-binding site are provided in Table 2C.

**Table 2.** (C): Active site and metal binding pocket analysis of *Clostridial AAD* enzymes

| Enzyme | Putative active site residues | Putative metal binding site | Other residues | Catalytic residue |
| --- | --- | --- | --- | --- |
| *adhE1/E2* | D485-G543-T544-P545-S548-K551-T5595-S596-G598-K617-Y618-L636-L644-D651-N655 | D651-N655-H721-S735 | D651 (Zinc metal, catalytic)<br>D485 (NAD binding residue) | C244 |
| *bdh/A/B* | G39-G40-S41-P70-N71-G97-G98-S99-D102-T138-L139-T142-S144-D147-I149-V151-K160-Y179-T182-V183-D194-H198-H267-H281 | D194-H198-H267-H281 | Dimer interface: P9-K11-V12-F14-G15-R222-K226-A242-N243-W246 | S140 |
| *Aydh* | D486-G544-G545-S546-D549-K552-T596-S597-G599-K618-Y619-L637-L645-D652-H656-H722-H726-H736- | D652-H656-H722-H736 | - | C244 |
| *gap3dh* | A148-S149-C150-T151-T152-N153-C154-L155 | - | NAD binding: R11-I12, D33, S119, N314<br>G3P binding: S149-C150-T151, T210-G211, T180, T210-G211, R233 | C150, H177 |

### 3.4. DFT simulations-based generation and optimization of ligand structures

As mentioned in the subsection 2.2.2 above, the chemical structures of 17 ligands comprising 10 FI and 7 representative substrates were designed in GaussView v6.0 software and geometrically optimized in Gaussian16. All geometrically DFT-optimized ligand structure was then subjected for energy minimization. The final ligand structures used for molecular docking simulation are provided in Fig. 2B.

**Fig. 2.**
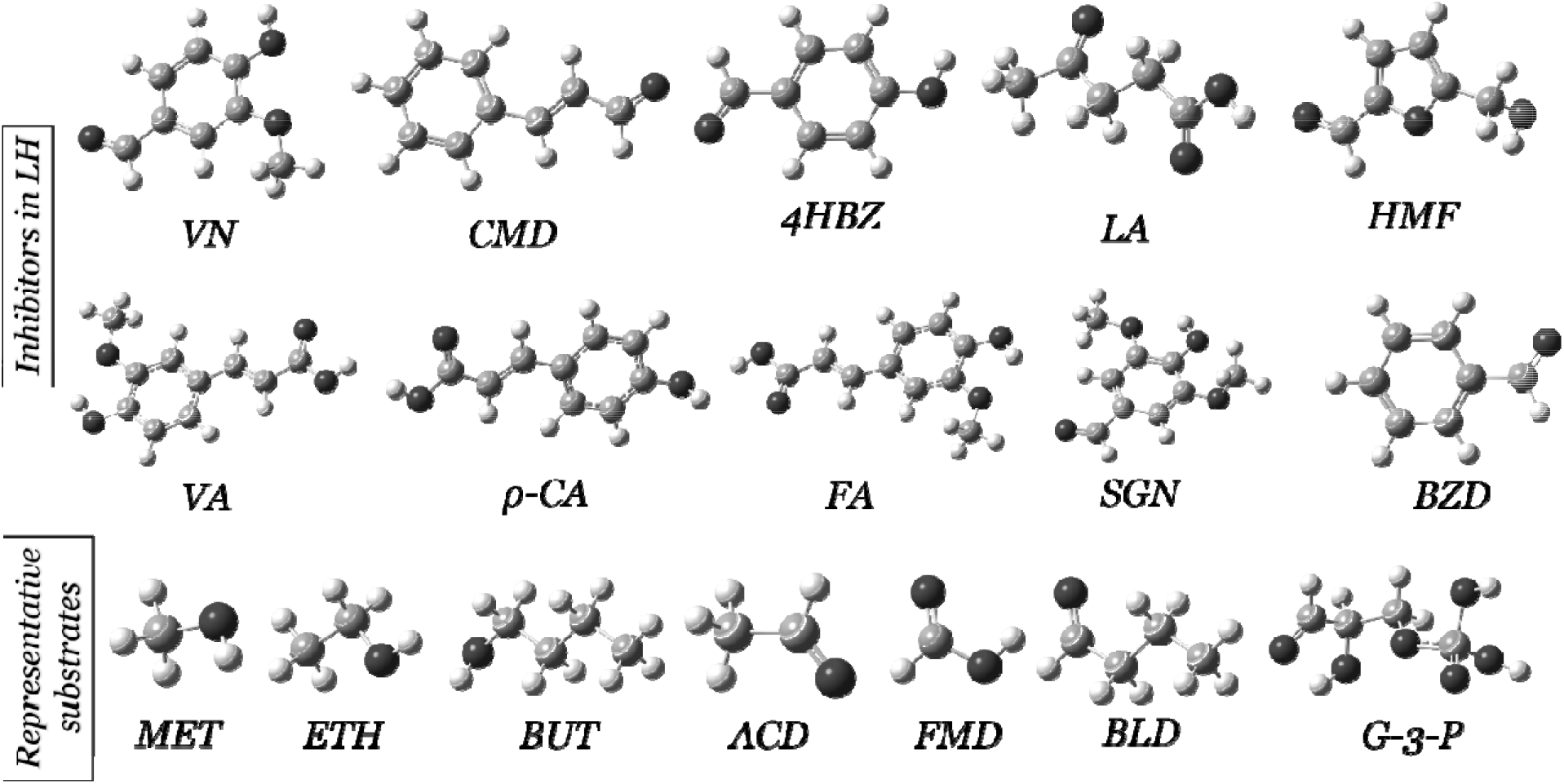
(B): The final 3-dimensional structure of DFT-based geometrically optimized and energy minimized ligands (fermentative inhibitors and representative substrates) used for molecular docking. VN – vanillin, CMD – cinnamaldehyde, 4HBZ – 4-hydroxy-benzaldehyde, LA – levulinic acid, HMF – 5-hyroxymethylfurfural, VA – vanillic acid, *ρ*CA – *ρ*-coumaric acid, FA – ferulic acid, SGN – syringaldehyd**e,** BZD – benzaldehyde, MET – methanol, ETH – ethanol, BUT – butanol, ACD – acetaldehyde, FMD – formaldehyde, BLD – butyraldehyde, and G-3-P – glyceraldehyde-3-phosphate

### 3.5. Molecular docking simulations of ligands with *Clostridial AAD* enzymes

The molecular docking of *Clostridial AAD* enzymes (Fig. 2A) with 17 ligands (Fig. 2B) is explained in detail in section 2.4. The molecular docking analysis revealed the binding energies (ΔG, kcal mol^-1^), inhibition constants (*K_i_*), and interacting residues with corresponding ligands. The binding energies for these 105 docking studies are summarised in Table 3A, where the sign and numerical value corresponds to extent and strength of binding with enzymes, respectively (negative sign implies favourable binding, and lower magnitude implies higher binding affinity). From Table 3A, it is clear that CMD and VA had highest binding affinity (i.e. lowest ΔG) with bifunctional enzymes, *adhE1* and *adhE2*. In addition to CMD and VA, *ρ*-CA showed highest binding affinity with aldehyde specific enzymes, *aydh* and *gap3dh*. In case of alcohol specific enzymes (*bdh, bdhA*, and *bdhB*) *ρ*-CA showed highest binding affinity. These selections of the potent inhibitor were made in comparison with the respective representative substrate (Fig. 2B). The selections of *ρ*-CA, CMD, and VA as the potent inhibitors are also supported by the lowest inhibition constant (Table 3B) as compared to other compounds in respective category. The results of docking analysis (i.e. interacting residues, type of interactions and nature of binding) of most potent inhibitors (*ρ*-CA, CMD, and VA) are summarised in Table 3C. The 3D and 2D visualization of the docked complex of *ρ*-CA, CMD, and VA are shown in Figs. 3(C, D, G, H, K, L, O and P).

**Fig. 3:**
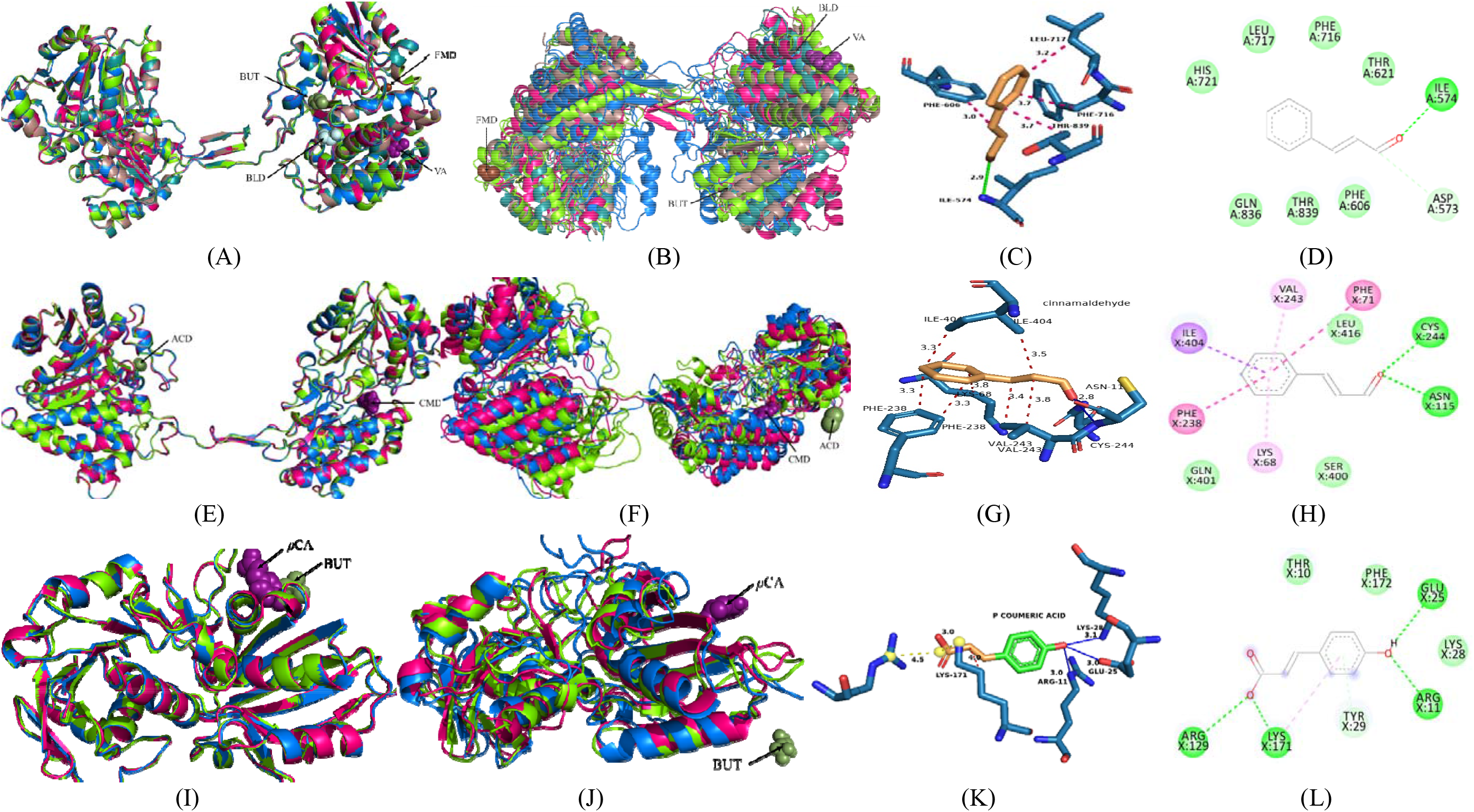
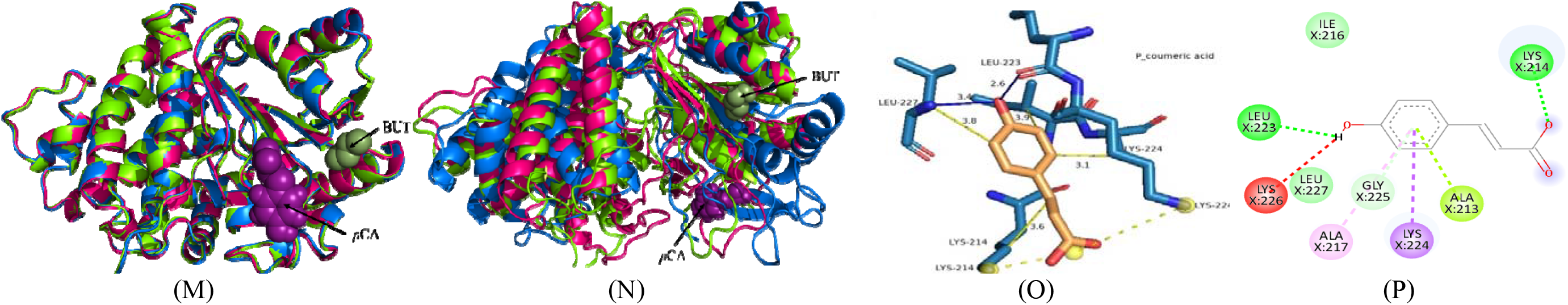
Analysis of comparative molecular docking and dynamics simulations of ligand-bound *AADs*. (A, E, I, M) – Superimposed initial ligand-bound structures of *AADs* before MD simulations, and (B, F, J, N) – Superimposed ligand-bound structures of *AADs* after MD simulations from categories *adhE1/E2, aydh, gap3dh, bdh/bdhA/bdhB*, respectively. The 3D and 2D representations of inhibitor-bound *AAD* structures visualized in PyMOL and Biovia discovery studio after molecular docking simulations: (C, D) – VA bound *adhE1/E2*, (G, H) – CMD bound *aydh*, (K, L) – *ρ*CA bound *gap3dh*, (O, P) – *ρ*-CA bound *bdh/bdhA/bdhB*, respectively.

**Table 3.** (A): Binding energy (ΔG, kcal mol^-1^) of 17 ligands with *Clostridial AAD* enzymes

| Ligands | Binding energy ($\Delta G$, kcal mol<sup>-1</sup>) | | | | | | |
| --- | --- | --- | --- | --- | --- | --- | --- |
|  | *adhE1* | *adhE2* | *bdhA* | *bdhB* | *bdh* | *aydh* | *gap3dh* |
| *Inhibitors* |  |  |  |  |  |  |  |
| Vanillin (VN) | -4.75 | -4.71 | -5.44 | -4.81 | -5.05 | -5.34 | -5.04 |
| Cinnamaldehyde (CMD) | -4.84 | -5.37 | -5.1 | -5.47 | -5.12 | -5.44 | -5.4 |
| 4-Hydroxy benzaldehyde (4HBZ) | -4.07 | -4.86 | -5.25 | -4.77 | -4.82 | -4.66 | -3.54 |
| Levulinic acid (LA) | -3.56 | -4.94 | -4.42 | -4.67 | -3.95 | -4.4 | -4.01 |
| 5-Hydroxymethylfurfural (HMF) | -3.63 | -4.57 | -4.63 | -4.59 | -4.45 | -4.55 | -5.13 |
| Vanillic acid (VA) | -4.89 | -5.17 | -5.76 | -5.75 | -5.18 | -5.12 | -5.8 |
| *p*-Coumeric acid (*p*-CA) | -4.36 | -5.15 | -5.97 | -5.88 | -5.46 | -5.23 | -6.19 |
| Ferulic acid (FA) | -3.92 | -5.11 | -5.76 | -5.7 | -4.98 | -5.19 | -5.49 |
| Syringaldehyde (SGN) | -4.54 | -4.83 | -4.79 | -4.94 | -5.17 | -3.69 | -5.13 |
| Benzaldehyde (BZD) | -4.07 | -4.61 | -4.95 | -4.41 | -4.58 | -4.55 | -4.65 |
| *Representative substrates* |  |  |  |  |  |  |  |
| Methanol (MET) | -2.18 | -2.36 | -2.88 | -2.55 | -2.3 | -2.6 | -1.92 |
| Ethanol (ETH) | -2.66 | -3.02 | -3.28 | -2.76 | -2.68 | -2.58 | -2.37 |
| Butanol (BUT) | -2.64 | -3.04 | -3.25 | -3.43 | -3.38 | NA | NA |
| Formaldehyde (FMD) | -3.34 | -2.86 | -3.13 | -4.02 | -3.14 | -3.35 | -3.05 |
| Acetaldehyde (ACD) | NA | NA | NA | NA | NA | -2.85 | NA |
| Butyraldehyde (BLD) | -3.22 | -3.52 | -3.39 | -3.46 | -3.17 | -2.6 | -2.62 |
| Glyceraldehyde-3-phosphate (G3P) | NA | NA | NA | NA | NA | NA | -3.65 |

**Table 3.** (B): System definitions for docking and MD simulations of selected ligands

| S. No. | Docking index | Receptor | Ligand | $K_i$ (mM) | Complex index for MD simulation | SASA (nm<sup>2</sup>) | RMSD (nm) |
| --- | --- | --- | --- | --- | --- | --- | --- |
| 1 | A0.pdb | *adhE1* | No ligand | NA | Control 0 = C0.gro (*adhE1* + water) | 389.9 | 1.03 |
| 2 | A1.pdb | *adhE1* | Butanol (BUT) | 11.6 | Test 1 = C1.gro (*adhE1* + BUT) | 395.5 | 1.01 |
| 3 | A2.pdb | *adhE1* | Formaldehyde (FMD) | 3.55 | Test 2 = C2.gro (*adhE1* + FMD) | 401.3 | 1.12 |
| 4 | A3.pdb | *adhE1* | Butyraldehyde (BLD) | 4.38 | Test 3 = C3.gro (*adhE1* + BLD) | 393.1 | 1.05 |
| 5 | A4.pdb | *adhE1* | Vanillic acid (VA) | 0.26 | Test 4 = C4.gro (*adhE1* + VA) | 391.4 | 0.81 |
| 6 | A5.pdb | *adhE2* | No ligand | NA | Control 1 = C5.gro (*adhE2* + water) | 402.4 | 1.04 |
| 7 | A6.pdb | *adhE2* | Butanol (BUT) | 5.96 | Test 5 = C6.gro (*adhE2* + BUT) | 400.5 | 1.1 |
| 8 | A7.pdb | *adhE2* | Butyraldehyde (BLD) | 2.64 | Test 6 = C7.gro (*adhE2* + BLD) | 397.1 | 1.33 |
| 9 | A8.pdb | *adhE2* | Cinnamaldehyde (CMD) | 0.12 | Test 7 = C8.gro (*adhE2* + CMD) | 401.8 | 0.98 |
| 10 | B0.pdb | *bdh* | No ligand | NA | Control 2 = C9.gro (*bdh* + water) | 183.4 | 0.29 |
| 11 | B1.pdb | *bdh* | Butanol (BUT) | 3.30 | Test 8 = C10.gro (*bdh* + BUT) | 186.5 | 0.26 |
| 12 | B2.pdb | *bdh* | ρ-Coumeric acid (ρ-CA) | 0.1 | Test 9 = C11.gro (*bdh* + ρ-CA) | 187.2 | 0.28 |
| 13 | B3.pdb | *bdhA* | No ligand | NA | Control 3 = C12.gro (*bdhA* + water) | 187.5 | 0.34 |
| 14 | B4.pdb | *bdhA* | Butanol (BUT) | 4.14 | Test 10 = C13.gro (*bdhA* + BUT) | 182.7 | 0.36 |
| 15 | B5.pdb | *bdhA* | ρ-Coumeric acid (ρ-CA) | 0.05 | Test 11 = C14.gro (*bdhA* + ρ-CA) | 187.3 | 0.33 |
| 16 | B6.pdb | *bdhB* | No ligand | NA | Control 4 = C15.gro (*bdhB* + water) | 196.8 | 0.80 |
| 17 | B7.pdb | *bdhB* | Butanol (BUT) | 3.08 | Test 12 = C16.gro (*bdhB* + BUT) | 189.8 | 0.35 |
| 18 | B8.pdb | *bdhB* | ρ-Coumeric acid (ρ-CA) | 0.05 | Test 13 = C17.gro (*bdhB* + ρ-CA) | 213.3 | 0.40 |
| 19 | D0.pdb | *aydh* | No ligand | NA | Control 5 = C18.gro (*aydh* + water) | 399.5 | 0.94 |
| 20 | D1.pdb | *aydh* | Acetaldehyde (ACD) | 8.21 | Test 14 = C19.gro (*aydh* + ACD) | 401.7 | 1.26 |
| 21 | D2.pdb | *aydh* | Cinnamaldehyde (CMD) | 0.1 | Test 15 = C20.gro (*aydh* + CMD) | 404.5 | 1.14 |
| 22 | D3.pdb | *gap3dh* | No ligand | NA | Control 6 = C21.gro (*gap3dh* + water) | 169.2 | 0.28 |
| 23 | D4.pdb | *gap3dh* | Glyceraldehyde-3-phosphate (G3P) | 2.10 | Test 16 = C22.gro (*gap3dh* + G3P) | 166.0 | 0.33 |
| 24 | D5.pdb | *gap3dh* | ρ-Coumeric acid (ρ-CA) | 0.03 | Test 17 = C23.gro (*gap3dh* + ρ-CA) | 168.9 | 0.26 |

**Abbreviation:** $K_i$ – inhibition constant; MD – molecular dynamics; SASA – solvent accessible surface area; RMSD – root mean square deviation

**Table 3.**
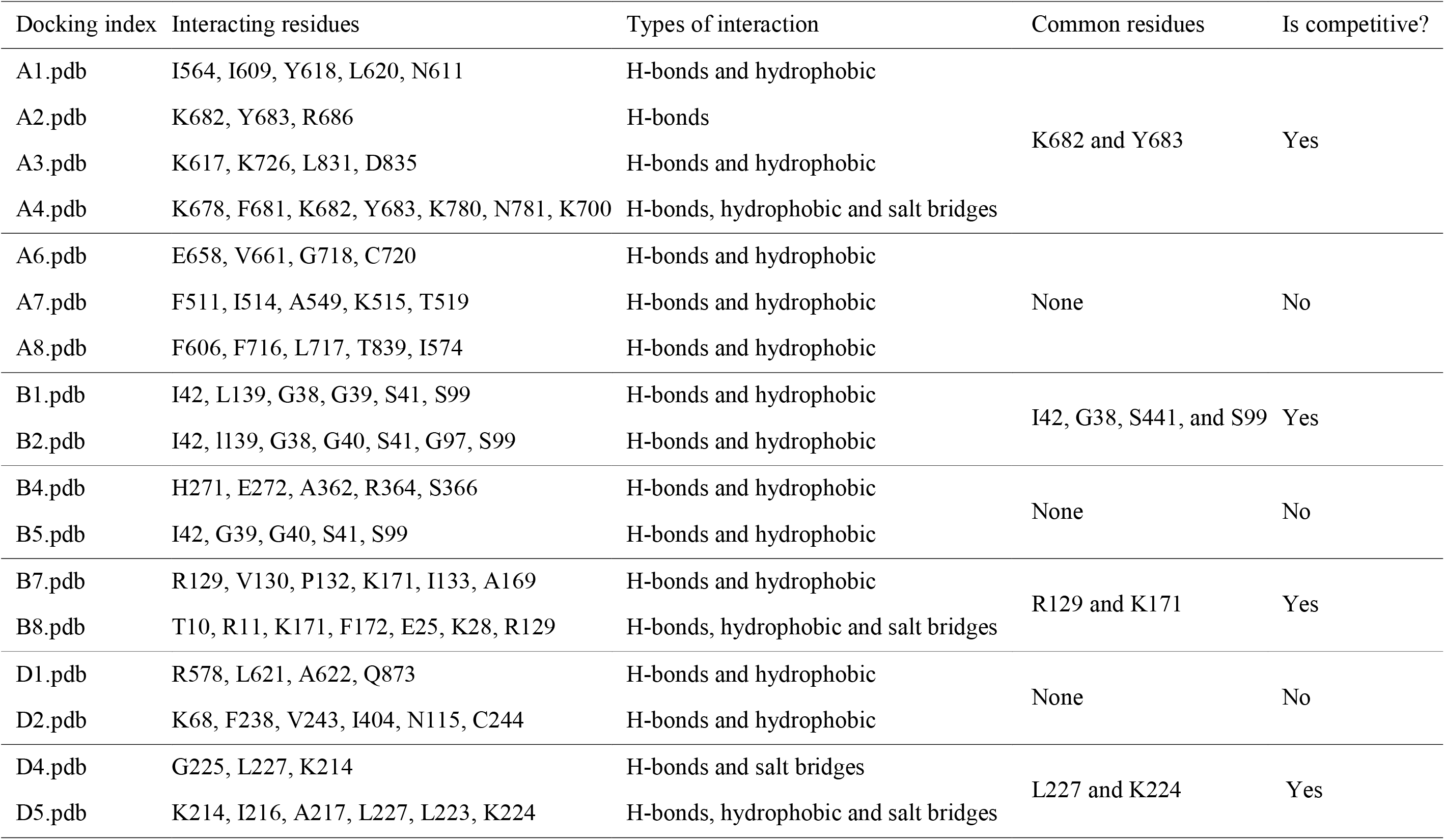
C): Comparative molecular docking analysis to deduce type of inhibition by VA, CMD, and *ρ*-CA

### 3.6. Comparative MD Analysis of ligand-bound *AAD* trajectories

The best docking complexes of each *AAD* enzyme (including representative substrates and inhibitors), listed in Table 3B, were tested further for its destabilization potential by means of MD simulation. The definition of all such docked complexes and MD simulation systems subjected for dynamic analysis are provided in Table 3B. The MD simulation studies were conducted for 50 ns for all 24 MD systems (7 control and 17 test systems) under investigation (see Table 3B). The analysis of comparative MD simulation of ligand-bound *AADs* are shown in Figs. 3(A, B, E, F, I, J, M, N) where Figs. 3(A, E, I, M) represents superimposed conformation of ligand-bound initial (i.e. 0 ns) gromacs structures and Figs. 3(B, F, J, N) represents superimposed conformation of ligand-bound final (i.e. after 50 ns) gromacs structures of *AADs* from category *adhE1/E2, aydh, gap3dh*, and *bdh/bdhA/bdhB*. The *ρ*-CA, CMD, and VA (the strongest inhibitors) were observed to migrate to different probable binding sites along the *AAD* enzymes but the average ensemble image has been depicted for all the ligands under investigation in Figs. 3(A, B, E, F, I, J, M, N). A clear demarcation of the disorganization in final *AADs* gromacs structures was observed in the presence of *ρ*-CA, CMD, and VA with respect to representative substrates BUT, BLD, ACD, and G3P (Table 3B), which imply the strong disrupting potential of these three FIs. Average RMSD and SASA over the 50 ns simulated trajectories are for all 24 MD systems are summarised in Table 3(B). The highest percentage change in average RMSD values were observed in the case of *ρ*-CA bound *bdhB* (50 % reduction) and VA bound *adhE1* (21 % reduction) as compared to the representative substrate, viz. BUT, FMD, and BLD (Table 3B). These reduction in the RMSD value are concurrent with rise in SASA which implies that *ρ*-CA and VA have higher disrupting potential by altering the backbone and impose structural instability of *adhE1* and *bdhB*. Similar trends (rise and reduction in RMSD with representative substrates and FIs, respectively) were also observed in case of other ligand-bound complexes in Table 3B. In conclusion, the primary analysis of investigating global MD trajectory of all 17 ligand-bound complex indicates *ρ*-CA as the most potent disruptor of the solventogenic *AADs*.

#### 3.6.1 Detailed comparative structural analysis of *AADs* in presence of *ρ*-CA, CMD, and VA

In addition to the calculation of average RMSD and SASA of the ligand-bound *AAD* structures, various other structural stability parameters such as Rg, RMSF, and H-bond formation were calculated for all 24 MD systems to screen the best disruptor for each *AADs* (explained in methodology section 2.5). Figs. 4 (A, B, C) shows the comparative Rg, H-bonds formation, and RMSF analysis, respectively, of MD simulations trajectory for ligand-bound *AADs* and *AADs* alone. These analyses are provided for the 4 different *AAD* categories, viz. *adhE1/E2, aydh, gap3dh*, and *bdh/bdhA/bdhB*.

**Fig 4:**
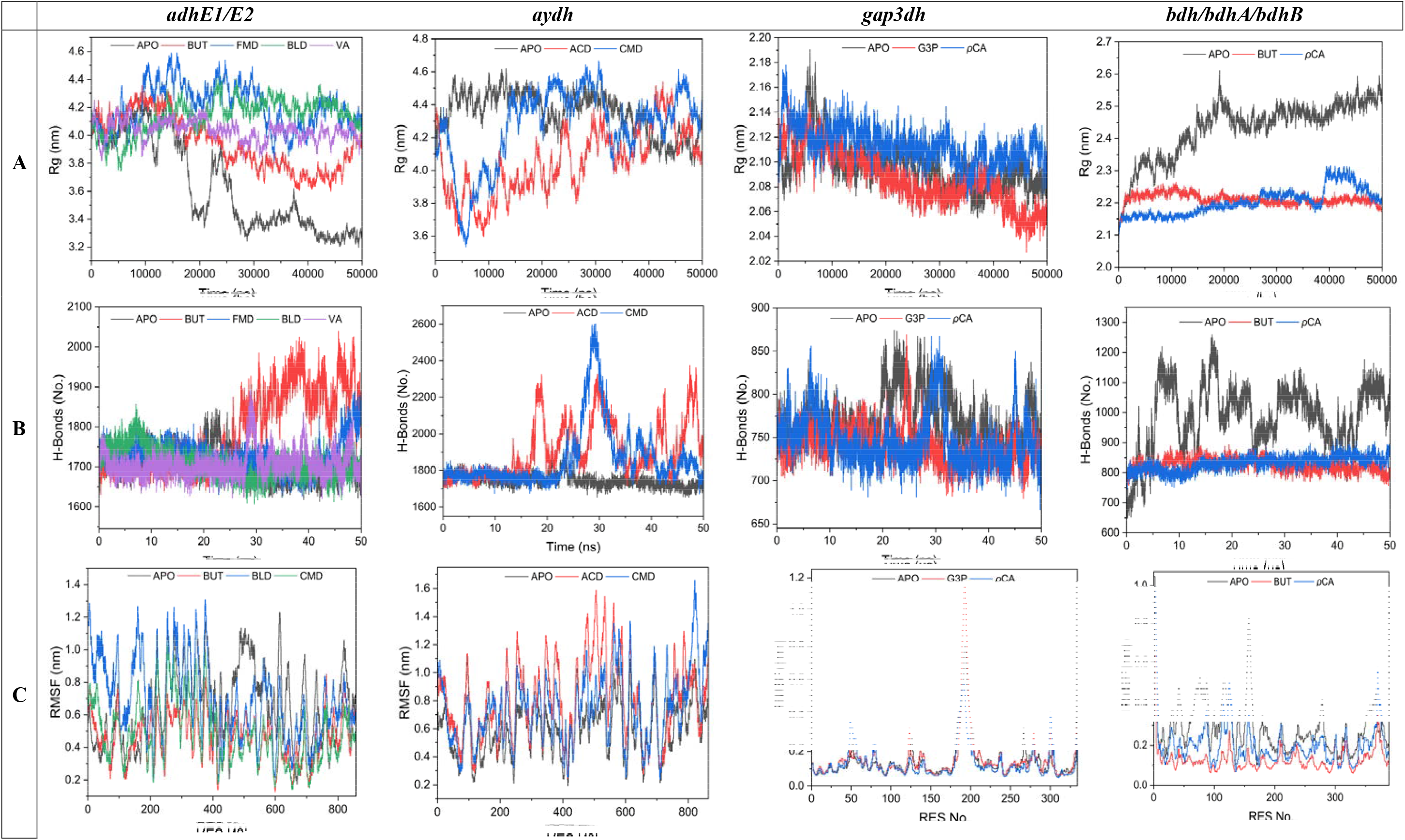
Analysis of comparative molecular dynamics simulations for ligand-bound *AADs* and *AADs* alone. (A) Rg analysis, (B) H-bonds analysis, (C) RMSF analysis for *AAD* categories *adhE1/E2, aydh, gap3dh*, and *bdh/bdhA/bdhB* enzymes.

## 4. Conclusions

Overall, our study conclud that *ρ*-CA is the strongest inhibitor since it shows highest binding affinities with 4 modelled enzymes. Furthermore, VA and CMD have also shown to inhibit 2 modelled enzymes each, indicating them to be the second strongest inhibitor. This analysis shows clearly that by minimizing lignocellulosic hydrolysate inhibitors specially *ρ*-CA, VA, and CMD, the enzymatic production of butanol in terms of yield, could enhance significantly while efficiently utilizing the biomass thus making ABE fermentation economically feasible. Moreover, this study provides a general framework which could applied for deciphering the molecular mechanism of inhibition behaviour of other enzymes in the metabolic pathway of solventogenic *Clostridia*.

## 5. Author contributions

KK: Conceptualization, Methodology, Formal Analysis, Validation, Software, Visualization, Investigation, Writing – Original draft, Review and Editing; ASMD, P, PC, MK., AS, and SY: Methodology, Formal Analysis, Investigation, Software, Visualization, Writing – Original draft; L.B.: Writing – Review; VSM: Supervision, Project administration, Writing – Review and Editing.

## Acknowledgments

Mr. Karan Kumar is grateful to the Prime Minister’s Research Fellowship provided by the Ministry of Education, Government of India. Authors acknowledge the fruitful discussions on analysis of results with Dr. Darpan Raghav, Biophysics and BioInnovations Laboratory, Indian Institute of Technology (IIT) Bombay. Use of HPC parallel computational facility, PARAM-ISHAN at IIT Guwahati is highly acknowledged. This research did not receive any specific grant from funding agencies in the public, commercial, or not–for–profit sectors.

## Reference

Agbor, V.B., Cicek, N., Sparling, R., Berlin, A., Levin, D.B., 2011. Biomass pretreatment: Fundamentals toward application. Biotechnology Advances 29, 675–685. https://doi.org/10.1016/j.biotechadv.2011.05.005

Baral, N.R., Shah, A., 2014. Microbial inhibitors: formation and effects on acetone-butanol-ethanol fermentation of lignocellulosic biomass. Appl Microbiol Biotechnol 98, 9151–9172. https://doi.org/10.1007/s00253-014-6106-8

Beg, Md.A., Shivangi, Thakur, S.C., Meena, L.S., 2018. Structural Prediction and Mutational Analysis of Rv3906c Gene of *Mycobacterium tuberculosis* H 37 Rv to Determine Its Essentiality in Survival. Advances in Bioinformatics 2018, 1–12. https://doi.org/10.1155/2018/6152014

Bhatia, S.K., Jagtap, S.S., Bedekar, A.A., Bhatia, R.K., Patel, A.K., Pant, D., Rajesh Banu, J., Rao, C.V., Kim, Y.-G., Yang, Y.-H., 2020. Recent developments in pretreatment technologies on lignocellulosic biomass: Effect of key parameters, technological improvements, and challenges. Bioresource Technology 300, 122724. https://doi.org/10.1016/j.biortech.2019.122724

Borah, A.J., Agarwal, M., Poudyal, M., Goyal, A., Moholkar, V.S., 2016. Mechanistic investigation in ultrasound induced enhancement of enzymatic hydrolysis of invasive biomass species. Bioresource Technology 213, 342–349. https://doi.org/10.1016/j.biortech.2016.02.024

Cao, G., Sheng, Y., 2016. Biobutanol Production from Lignocellulosic Biomass: Prospective and Challenges. J Bioremediat Biodegrad 7. https://doi.org/10.4172/2155-6199.1000363

Carrillo-Nieves, D., Rostro Alanís, M.J., de la Cruz Quiroz, R., Ruiz, H.A., Iqbal, H.M.N., Parra-Saldívar, R., 2019. Current status and future trends of bioethanol production from agroindustrial wastes in Mexico. Renewable and Sustainable Energy Reviews 102, 63–74. https://doi.org/10.1016/j.rser.2018.11.031

Chen, J.-S., 1995. Alcohol dehydrogenase: multiplicity and relatedness in the solvent-producing clostridia. FEMS Microbiol Rev 17, 263–273. https://doi.org/10.1111/j.1574-6976.1995.tb00210.x

Cho, C., Hong, S., Moon, H.G., Jang, Y.-S., Kim, D., Lee, S.Y., 2019. Engineering Clostridial Aldehyde/Alcohol Dehydrogenase for Selective Butanol Production. mBio. https://doi.org/10.1128/mBio.02683-18

Curtiss, L.A., Redfern, P.C., Raghavachari, K., 2007. Gaussian-4 theory. The Journal of Chemical Physics 126, 084108. https://doi.org/10.1063/1.2436888

Dahman, Y., 2012. Sustainable Biobutanol and Working towards the Green Gasoline of the Future. Fermentat Technol 01. https://doi.org/10.4172/2167-7972.1000e111

Dai, Z., Dong, H., Zhang, Y., Li, Y., 2016. Elucidating the contributions of multiple aldehyde/alcohol dehydrogenases to butanol and ethanol production in Clostridium acetobutylicum. Sci Rep 6, 28189. https://doi.org/10.1038/srep28189

Detain, J., Rémond, C., Rodrigues, C.M., Harakat, D., Besaury, L., 2022. Co-elicitation of lignocelluloytic enzymatic activities and metabolites production in an Aspergillus-Streptomyces co-culture during lignocellulose fractionation. Current Research in Microbial Sciences 3, 100108. https://doi.org/10.1016/j.crmicr.2022.100108

Fatma, S., Hameed, A., Noman, M., Ahmed, T., Shahid, M., Tariq, M., Sohail, I., Tabassum, R., 2018. Lignocellulosic Biomass: A Sustainable Bioenergy Source for the Future. PPL 25, 148–163. https://doi.org/10.2174/0929866525666180122144504

Gabler, F., Nam, S., Till, S., Mirdita, M., Steinegger, M., Söding, J., Lupas, A.N., Alva, V., 2020. Protein Sequence Analysis Using the MPI Bioinformatics Toolkit. Current Protocols in Bioinformatics 72. https://doi.org/10.1002/cpbi.108

Gaussian 09 Citation | Gaussian.com [WWW Document], n.d. URL https://gaussian.com/g09citation/ (accessed 4.17.21).

Guex, N., Peitsch, M.C., 1997. SWISS-MODEL and the Swiss-Pdb Viewer: An environment for comparative protein modeling. Electrophoresis 18, 2714–2723. https://doi.org/10.1002/elps.1150181505

Heo, L., Park, H., Seok, C., 2013. GalaxyRefine: protein structure refinement driven by side-chain repacking. Nucleic Acids Research 41, W384–W388. https://doi.org/10.1093/nar/gkt458

Jendele, L., Krivak, R., Skoda, P., Novotny, M., Hoksza, D., 2019. PrankWeb: a web server for ligand binding site prediction and visualization. Nucleic Acids Research 47, W345–W349. https://doi.org/10.1093/nar/gkz424

Jönsson, L.J., Martín, C., 2016. Pretreatment of lignocellulose: Formation of inhibitory by-products and strategies for minimizing their effects. Bioresource Technology 199, 103–112. https://doi.org/10.1016/j.biortech.2015.10.009

Kim, D., 2018. Physico-Chemical Conversion of Lignocellulose: Inhibitor Effects and Detoxification Strategies: A Mini Review. Molecules 23, 309. https://doi.org/10.3390/molecules23020309

Kim, Y., Ximenes, E., Mosier, N.S., Ladisch, M.R., 2011. Soluble inhibitors/deactivators of cellulase enzymes from lignocellulosic biomass. Enzyme and Microbial Technology 48, 408–415. https://doi.org/10.1016/j.enzmictec.2011.01.007

Kolesinska, B., Fraczyk, J., Binczarski, M., Modelska, M., Berlowska, J., Dziugan, P., Antolak, H., Kaminski, Z.J., Witonska, I.A., Kregiel, D., 2019. Butanol Synthesis Routes for Biofuel Production: Trends and Perspectives. Materials (Basel) 12, E350. https://doi.org/10.3390/ma12030350

Kour, D., Rana, K.L., Yadav, N., Yadav, A.N., Rastegari, A.A., Singh, C., Negi, P., Singh, K., Saxena, A.K., 2019. Technologies for Biofuel Production: Current Development, Challenges, and Future Prospects, in: Rastegari, A.A., Yadav, A.N., Gupta, A. (Eds.), Prospects of Renewable Bioprocessing in Future Energy Systems, Biofuel and Biorefinery Technologies. Springer International Publishing, Cham, pp. 1–50. https://doi.org/10.1007/978-3-030-14463-0_1

Kumar, K., Patro, P., Raut, U., Yadav, V., Barbora, L., Moholkar, V.S., 2022a. Ultrasonic enhancement of lipase-catalyzed reactions: Mechanistic investigation using molecular docking analysis (preprint). Preprints. https://doi.org/10.22541/au.166722851.12958171/v1

Kumar, K., Roy, K., Moholkar, V.S., 2021. Mechanistic investigations in sonoenzymatic synthesis of n-butyl levulinate. Process Biochemistry 111, 147–158. https://doi.org/10.1016/j.procbio.2021.09.005

Kumar, K., Shah, H., Moholkar, V.S., 2022b. Genetic Algorithm for Optimization of Fermentation Processes of Various Enzyme Productions, in: Optimization of Sustainable Enzymes Production. Chapman and Hall/CRC.

Laskowski, R.A., Swindells, M.B., 2011. LigPlot+: Multiple Ligand–Protein Interaction Diagrams for Drug Discovery. J. Chem. Inf. Model. 51, 2778–2786. https://doi.org/10.1021/ci200227u

Laskowski, RomanA., Rullmann, J.AntoonC., MacArthur, MalcolmW., Kaptein, R., Thornton, JanetM., 1996. AQUA and PROCHECK-NMR: Programs for checking the quality of protein structures solved by NMR. J Biomol NMR 8. https://doi.org/10.1007/BF00228148

Malani, R.S., Umriwad, S.B., Kumar, K., Goyal, A., Moholkar, V.S., 2019. Ultrasound–assisted enzymatic biodiesel production using blended feedstock of non–edible oils: Kinetic analysis. Energy Conversion and Management 188, 142–150. https://doi.org/10.1016/j.enconman.2019.03.052

Mayank, R., Ranjan, A., Moholkar, V.S., 2013. Mathematical models of ABE fermentation: review and analysis. Critical Reviews in Biotechnology 33, 419–447. https://doi.org/10.3109/07388551.2012.726208

Morris, G.M., Huey, R., Lindstrom, W., Sanner, M.F., Belew, R.K., Goodsell, D.S., Olson, A.J., 2009. AutoDock4 and AutoDockTools4: Automated docking with selective receptor flexibility. J. Comput. Chem. 30, 2785–2791. https://doi.org/10.1002/jcc.21256

Morris, G.M., Huey, R., Olson, A.J., 2008. Using AutoDock for Ligand Receptor Docking. Current Protocols in Bioinformatics 24. https://doi.org/10.1002/0471250953.bi0814s24

Palmqvist, E., Hahn-Hägerdal, B., 2000. Fermentation of lignocellulosic hydrolysates. II: inhibitors and mechanisms of inhibition. Bioresource Technology 74, 25–33. https://doi.org/10.1016/S0960-8524(99)00161-3

Ranjan, A., Moholkar, V.S., 2012. Biobutanol: science, engineering, and economics: Review Essay. Int. J. Energy Res. 36, 277–323. https://doi.org/10.1002/er.1948

Rathour, R.K., Ahuja, V., Bhatia, R.K., Bhatt, A.K., 2018. Biobutanol: New era of biofuels. Int J Energy Res 42, 4532–4545. https://doi.org/10.1002/er.4180

Roy, A., Kucukural, A., Zhang, Y., 2010. I-TASSER: a unified platform for automated protein structure and function prediction. Nat Protoc 5, 725–738. https://doi.org/10.1038/nprot.2010.5

Singh, N., Kumar, K., Goyal, A., Moholkar, V.S., 2022. Ultrasound-assisted biodiesel synthesis by in–situ transesterification of microalgal biomass: Optimization and kinetic analysis. Algal Research 61, 102582. https://doi.org/10.1016/j.algal.2021.102582

Singhvi, M.S., Chaudhari, S., Gokhale, D.V., 2014. Lignocellulose processing: a current challenge. RSC Adv. 4, 8271. https://doi.org/10.1039/c3ra46112b

Wallner, B., 2006. Identification of correct regions in protein models using structural, alignment, and consensus information. Protein Science 15, 900–913. https://doi.org/10.1110/ps.051799606

Yang, Y., Hu, M., Tang, Y., Geng, B., Qiu, M., He, Q., Chen, S., Wang, X., Yang, S., 2018. Progress and perspective on lignocellulosic hydrolysate inhibitor tolerance improvement in Zymomonas mobilis. Bioresources and Bioprocessing 5, 6. https://doi.org/10.1186/s40643-018-0193-9

Yoo, M., Croux, C., Meynial-Salles, I., Soucaille, P., 2016. Elucidation of the roles of adhE1 and adhE2 in the primary metabolism of Clostridium acetobutylicum by combining in-frame gene deletion and a quantitative system-scale approach. Biotechnol Biofuels 9, 92. https://doi.org/10.1186/s13068-016-0507-0

Yuan, S., Chan, H.C.S., Hu, Z., 2017. Using PYMOL as a platform for computational drug design. WIREs Comput Mol Sci 7. https://doi.org/10.1002/wcms.1298

Zimmermann, L., Stephens, A., Nam, S.-Z., Rau, D., Kübler, J., Lozajic, M., Gabler, F., Söding, J., Lupas, A.N., Alva, V., 2018. A Completely Reimplemented MPI Bioinformatics Toolkit with a New HHpred Server at its Core. Journal of Molecular Biology 430, 2237–2243. https://doi.org/10.1016/j.jmb.2017.12.007

